# Comprehensive evaluation of computational cell-type quantification methods for immuno-oncology

**DOI:** 10.1101/463828

**Authors:** Gregor Sturm, Francesca Finotello, Florent Petitprez, Jitao David Zhang, Jan Baumbach, Wolf H. Fridman, Markus List, Tatsiana Aneichyk

## Abstract

**Motivation:** The composition and density of immune cells in the tumor microenvironment profoundly influence tumor progression and success of anti-cancer therapies. Flow cytometry, immunohistochemistry staining, or single-cell sequencing is often unavailable such that we rely on computational methods to estimate the immune-cell composition from bulk RNA-sequencing (RNA-seq) data. Various methods have been proposed recently, yet their capabilities and limitations have not been evaluated systematically. A general guideline leading the research community through cell type deconvolution is missing.

**Results:** We developed a systematic approach for benchmarking such computational methods and assessed the accuracy of tools at estimating nine different immune- and stromal cells from bulk RNA-seq samples. We used a single-cell RNA-seq dataset of ∼11,000 cells from the tumor microenvironment to simulate bulk samples of known cell type proportions, and validated the results using independent, publicly available gold-standard estimates. This allowed us to analyze and condense the results of more than a hundred thousand predictions to provide an exhaustive evaluation across seven computational methods over nine cell types and ∼1,800 samples from five simulated and real-world datasets. We demonstrate that computational deconvolution performs at high accuracy for well-defined cell-type signatures and propose how fuzzy cell-type signatures can be improved. We suggest that future efforts should be dedicated to refining cell population definitions and finding reliable signatures.

**Availability:** A snakemake pipeline to reproduce the benchmark is available at https://github.com/grst/immune_deconvolution_benchmark. An R package allows the community to perform integrated deconvolution using different methods (https://grst.github.io/immunedeconv).

## 1 Introduction

Tumors are not only composed of malignant cells but are embedded in a complex microenvironment within which dynamic interactions are built (Fridman *et al.* (2012)). Notably, this tumor microenvironment (TME) comprises a vast variety of immune cells. Knowledge of immune cell content in cancer samples is invaluable for cancer-immunotherapy drug discovery as well as for clinical decisions about treatment options. The cellular composition of the immune infiltrate of tumors can shed light on the escape mechanisms that tumor cells use to evade the immune response. In clinical trials, it can be used to stratify patients to assign most suitable treatment options depending on the targeted cell type, hence increasing the overall chances of success, and ultimately accelerating access to improved treatment options (Friedman *et al.* (2015)).

Methods like fluorescence-activated cell sorting (FACS) or immunohistochemistry (IHC)-staining have been used as gold standards to estimate the immune cell content within a sample (Petitprez *et al.* (2018a)). However, each of the methods has its technical limitations and might not be generally applicable. More specifically, FACS requires a large amount of material, hence limiting its application to tumor biopsies, and IHC provides an estimate from a single tumor slice, which may not be representative of a heterogeneous immune landscape of the tumor. Furthermore, both methods can utilize only a small number of cell type-specific markers. More recently, single-cell RNA sequencing (scRNA-seq) is used to characterize cell types and states, yet for the time being it remains too expensive and laborious for routine clinical use. Moreover, cell type proportions might be biased in scRNA-seq data due to differences in single-cell dissociation efficiencies (Lambrechts *et al.* (2018)). At the same time, methods for gene expression profiling, RNA-seq, and microarrays, have been developed and optimized to be applicable in clinical settings, resulting in a plethora of transcriptomics datasets derived from patient tumor samples. However, while these methods provide transcriptomics data from heterogeneous samples considered as a whole, they do not provide information on their cellular composition, which therefore has to be inferred using computational techniques.

Previously, available methods for estimating immune cell contents of tumor samples have been reviewed in terms of methodology (Finotello and Trajanoski (2018); Newman and Alizadeh (2016); Avila Cobos *et al.* (2018); Petitprez *et al.* (2018a)) and limited quantitative comparisons have been made as part of some of the methods’ publications (Racle *et al.* (2017); Becht *et al.* (2016); Aran *et al.* (2017)). These are however limited to only a few methods, do not address background predictions or are not cell-type specific. We, therefore, provide the first systematic, quantitative comparison of computational cell type quantification methods for immuno-oncology at a per-cell-type resolution. While we acknowledge that many general-purpose deconvolution methods are available (reviewed in Finotello and Trajanoski (2018)), we focus this benchmark on methods that provide cell-type signatures for immuno-oncology (table 1).

**Table 1.** Overview of cell type quantification methods providing gene signatures for immuno-oncology. Methods can be conceptually distinguished in marker-gene-based approaches (M) and deconvolution-based approaches (D). The output scores of the methods have different properties and allow either intra-sample comparisons between cell types, inter-sample comparisons of the same cell type, or both. All methods come with a set of cell type signatures ranging from 6 immune cell types to 64 immune and non-immune cell types.

| tool | abbrev. | type | score | comparisons | algorithm | cell types | reference |
| --- | --- | --- | --- | --- | --- | --- | --- |
| CIBERSORT | CBS | D | immune cell fractions, relative to total immune cell content | intra | ν-support vector regression | 22 immune cell types | Newman *et al.* (2015) |
| CIBERSORT abs. mode | CBA | D | score of arbitrary units that reflects the absolute proportion of each cell type | intra, inter | ν-support vector regression | 22 immune cell types | Newman *et al.* (2015, 2018) |
| EPIC | EPC | D | cell fractions, relative to all cells in sample | intra, inter | constrained least square regression | 6 immune cell types, fibroblasts, endothelial cells | Racle *et al.* (2017) |
| MCP-counter | MCP | M | arbitrary units, comparable between samples | inter | mean of marker gene expression | 8 immune cell types, fibroblasts, endothelial cells | Becht *et al.* (2016) |
| quanTIseq | QTS | D | cell fractions, relative to all cells in sample | intra, inter | constrained least square regression | 10 immune cell types | Finotello *et al.* (2017) |
| TIMER | TMR | D | arbitrary units, comparable between samples (not different cancer types) | inter | linear least square regression | 6 immune cell types | Li *et al.* (2016) |
| xCell | XCL | M | arbitrary units, comparable between samples | inter | ssGSEA (Hänzelmann *et al.* (2013)) | 64 immune and non-immune cell types | Aran *et al.* (2017) |

These methods can, in general, be classified into two categories: marker gene-based approaches and deconvolution-based approaches. Marker gene-based approaches utilize a list of genes that are characteristic for a cell type. These gene sets are usually derived from targeted transcriptomics studies characterizing each immune-cell type and/or from comprehensive literature search and experimental validation. By using the expression values of marker genes in heterogeneous samples, these models quantify every cell type independently, either aggregating them into an abundance score (MCP-counter, Becht *et al.* (2016)) or by performing a statistical test for enrichment of the marker genes (xCell, Aran *et al.* (2017)). Deconvolution methods, on the other hand, formulate the problem as a system of equations that describe the gene expression of a sample as the weighted sum of the expression profiles of the admixed cell types (reviewed in Finotello and Trajanoski (2018)). By solving the inverse problem, cell type fractions can be inferred given a signature matrix and the bulk gene expression. In practice, this problem can be solved using linear least square regression (*e.g.* TIMER, Li *et al.* (2016)), constrained least square regression (*e.g.* quanTIseq and EPIC, Finotello *et al.* (2017); Racle *et al.* (2017)), or ν-Support Vector Regression (*e.g.* CIBERSORT, Newman *et al.* (2015)).

Conducting a fair benchmark between computational cell type quantification methods is challenging, as the scores of the different methods have different properties and are not always directly comparable (table 1). For instance, an inherent limitation of marker-based approaches is that they only allow generating a semi-quantitative score, which enables a comparison between samples, but not between cell types (Petitprez *et al.* (2018b)). Deconvolution allows, in principle, to generate absolute scores that can be interpreted as cell fractions and compared both inter- and intra-sample. In practice, CIBERSORT expresses results as a fraction relative to the immune-cell content and TIMER produces a score in arbitrary units that cannot be compared between cell types. Both quanTIseq and EPIC generate scores relative to the total amount of sequenced cells, being the only two methods generating an absolute score that can be interpreted as a cell fraction. CIBERSORT has recently been extended to the “absolute mode”, which provides a score that can be compared between both samples and cell types but still cannot be interpreted as a cell fraction (Newman *et al.* (2018)). From now on, we refer to this extension as “CIBERSORT abs”. quanTIseq comes as an entire pipeline that allows to directly analyze raw RNA-seq data (*i.e.* FASTQ files of sequencing reads), whereas the other methods only analyze pre-computed, normalized expression data. In this benchmark, we consider only its “deconvolution” module alone, acknowledging that the full quanTIseq pipeline can, in principle, provide higher performance. A limitation of TIMER and xCell is that the results of these methods depend on all samples that are analyzed in a single run, i.e. the same sample can have different scores when submitted together with different other samples. In particular, xCell uses the variability among the samples for a linear transformation of the output score. TIMER uses COMBAT (Johnson *et al.* (2007)) to merge the input samples with reference profiles. This is particularly problematic with small datasets and hampers comparability and interpretability of the score. Moreover, xCell does not detect any signal when ran with few non-heterogeneous samples (Aran (2018)). Nonetheless, all methods except CIBERSORT (relative mode) support between-sample comparisons and the performance of the methods can be readily compared using Pearson correlation with the gold-standard measure of cell fractions.

Here we developed a systematic approach for benchmarking such computational methods and assessed the accuracy of estimating the abundance of nine immune and stromal cell types from bulk RNA-seq samples for seven state-of-the-art approaches specifically designed for immunooncology and provide immuno-oncologists guidance on which method they can apply in their studies. We use a scRNA-seq dataset of more than 11,000 single cancer, stromal, and immune cells (Schelker *et al.* (2017)) to simulate bulk RNA-seq samples with known compositions and validate the benchmark results using three independent datasets that have been profiled with FACS (Schelker *et al.* (2017); Hoek *et al.* (2015); Racle *et al.* (2017)). We assess the performance of the methods by four metrics, namely: (i) predictive performance, (ii) minimal detection fraction, (iii) background predictions and (iv) spillover effects. We demonstrate, that by addressing spillover effects, ‘fuzzy’ signatures of still not well-understood cell-types can be significantly improved. Finally, we provide an R package, `immunedeconv`, integrating all of the investigated methods for user-friendly access. To facilitate benchmarking of newly developed methods and signatures, we provide a reproducible pipeline on GitHub, which can be straightforwardly extended to benchmark additional methods.

## 2 Results

### Benchmark on simulated and true bulk RNA-seq samples reveals differences in method performance between cell types

We created 100 simulated bulk RNA-seq samples with known cell type proportions using the single cell dataset. For each sample, we individually retrieved and aggregated 500 random immune- and non-immune cells. This approach has been established and successfully applied by Schelker *et al.* (2017) for benchmarking CIBERSORT. Additional consistency checks support that simulated bulk RNA-seq data is not subject to systematic biases (supplementary figures 1-4).

We applied the seven methods to these samples and compared the estimated to the known fractions. The results are shown in figure 1a. All methods obtained a high correlation on B cells (Pearson’s *r >* 0.71), cancer associated fibroblasts (*r >* 0.72), endothelial cells (*r >* 0.94) and CD8+ T cells (*r >* 0.76). Most methods obtain a correlation of *r >* 0.68 on macrophages/monocytes, NK cells and overall CD4+ T cells. We observed poor performance in distinguishing regulatory-from non-regulatory CD4+ T cells and in estimating dendritic cells (DCs).

We validated the results using three independent datasets, which provide both bulk RNA-seq data and a gold-standard estimate of immune cell fractions using FACS (figure 1b-d and supplementary figure 5). In general, the results are highly consistent with the mixing benchmark, with the exception that the methods’ performance on CD8+ T cells is worse on the validation data considered altogether (figure 1c). This can be attributed to the fact that the variance of CD8+ T cell contents between the samples is low and the correlations are therefore not meaningful (see figure 1b and supplementary figure 5). Moreover, in the simulation benchmark, only TIMER detects DCs, while in Hoek’s dataset (Hoek *et al.* (2015)), all methods except CIBERSORT detect a signal. This inconsistency, and the poor performance on DCs in general might be indicative of the still not well defined biological heterogeneity between DC subtypes.

**Figure 1.**
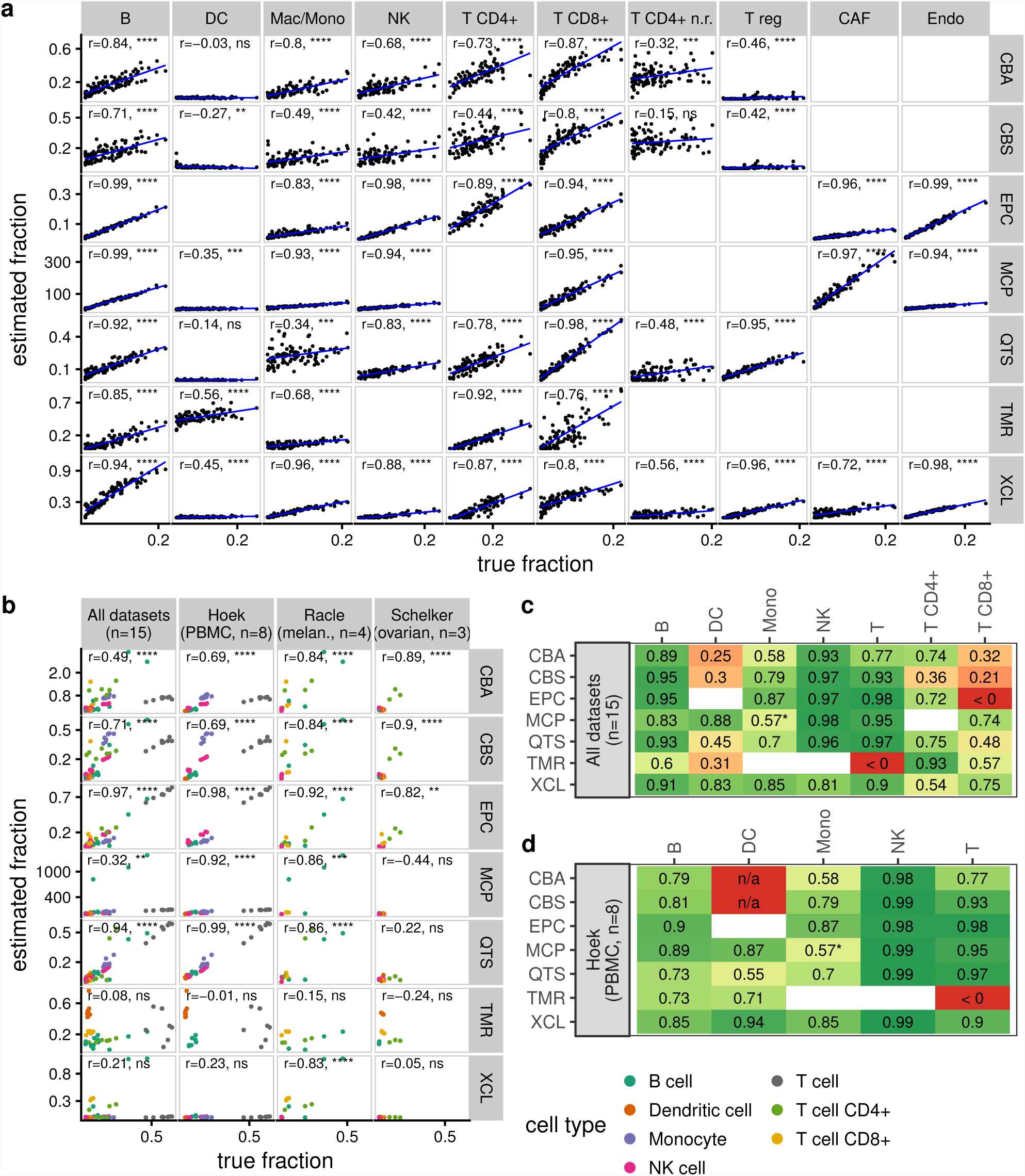
(a) Correlation of predicted vs. known cell type fractions on 100 simulated bulk RNA-seq samples generated from single cell RNA-seq. Pearson’s *r* is indicated in each panel. Due to the lack of a corresponding signature, we estimated macrophages/monocytes with EPIC using the “macrophage” signature and with MCP-counter using the “monocytic lineage” signature as a surrogate. (b) Performance of the methods on three independent datasets that provide immune cell quantification by FACS. Different cell types are indicated in different colors. Pearson’s *r* has been computed as a single correlation on all cell types simultaneously. Note that only methods that allow both inter- and intra-sample comparisons (i.e. EPIC, quanTIseq, CIBERSORT absolute mode) can be expected to perform well here. (c-d) Performance on the three validation datasets per cell type. Schelker’s and Racle’s dataset have too few samples to be considered individually. The values indicate Pearson correlation of the predictions with the cell type fractions determined using FACS. Blank squares indicate that the method does not provide a signature for the respective cell type. ‘n/a’ values indicate that no correlation could be computed because all predictions were zero. The asterisk (*) indicates that the ‘monocytic lineage’ signature was used as a surrogate to predict monocyte content. P-values: **** *<* 0.0001; *** *<* 0.001; ** *<* 0.01; * *<* 0.05; ns *≥* 0.05. P-values are not adjusted for multiple testing. Method abbreviations: see table 1.

### Background predictions widespread among deconvolution-based methods

Next we investigated two related questions: at which abundance level do methods reliably identify the presence of immune cells (minimal detection fraction) and, conversely, what fractions of a certain cell type are predicted even when they are actually absent (background predictions). To this end, we took advantage of the fact that single cell-based simulations allow us to create artificial bulk RNA-seq samples of arbitrary compositions. For each cell type, we created samples by spiking-in an increasing amount of the cell type of interest into a background of 1000 cells randomly sampled from all other cell types. We measured the background prediction level as the predicted score on the background only, *i.e.* no spike-in cells added. We defined the minimal detection fraction as the minimal number of spike-in cells needed for the score to be significantly different from the background.

We observed that, in most cases, the deconvolution-based approaches predict a minimal amount of immune cells even though they are absent (figure 2). Yet, background predictions are low for CAFs and NK cells for EPIC and non-regulatory CD4+ T cells and NK cells for quanTIseq. Also, TIMER does not suffer from background predictions (except for DCs) at the cost of a highly elevated minimal detection fraction. In contrast, xCell, which uses a statistical test for enrichment of the marker-genes, is highly robust against background-predictions (score = 0 for all tested cell types) at a slightly elevated minimal detection fraction (< 5 % infiltration for most cell types). Unfortunately, testing MCP-counter for its detection limit and background-predictions is not straightforward, as the method does not compute a score, but uses raw gene expression values. To make a statement about the absence of a cell type, a platform-specific null-model needs to be generated, which, in fact, is already provided by the authors for some microarray platforms, but not for RNA-seq. The authors also addressed the detection limit on RNA mixtures in their original publication (Becht *et al.* (2016)). In short, we observe that deconvolution-based approaches are susceptible to background-predictions that might be due to the similarity of the signatures of closely-related cell types (multicollinearity) and/or to a lower cell-specificity of signature/marker genes.

**Figure 2.**
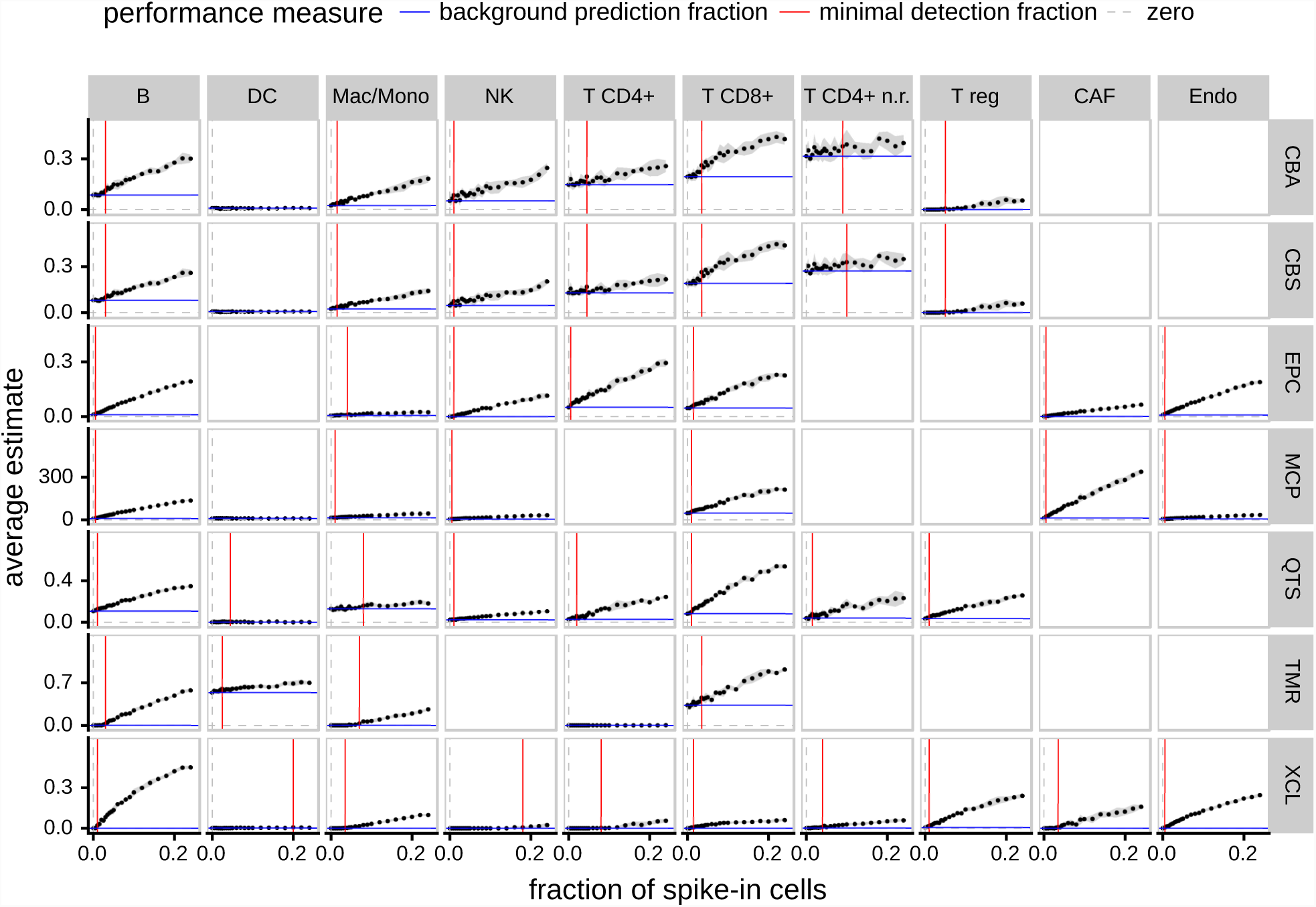
Minimal detection fraction and background prediction level. For each panel, we created simulated bulk RNA-seq samples with an increasing amount of the cell type of interest and a background of 1,000 cells randomly sampled from the other cell types. The dots show the mean predicted score across five independently simulated samples for each fraction of spike-in cells. The grey ribbon indicates the 95 % confidence interval. The red line refers to the minimal detection fraction, i.e. the minimal fraction of an immune cell type needed for a method to reliably detect its abundance as significantly different from the background (p-value < 0.05, one-sided t-test). The blue line refers to the background prediction level, i.e. the average estimate of a method while the cell type of interest is absent. Method abbreviations: see table 1.

### Spillover can be attributed to non-specific signature genes

Motivated by the hypothesis that background predictions might be due to non-specific signature genes, we asked which cell types lead to methods erroneously predicting a higher abundance of another. We refer to this effect as ‘spillover’. We assessed spillover effects using simulated bulk RNA-seq samples containing single immune cell types (figure 3) and validated the results using true bulk RNA-seq samples of FACS-purified immune cells (supplementary figure 6).

**Figure 3.**
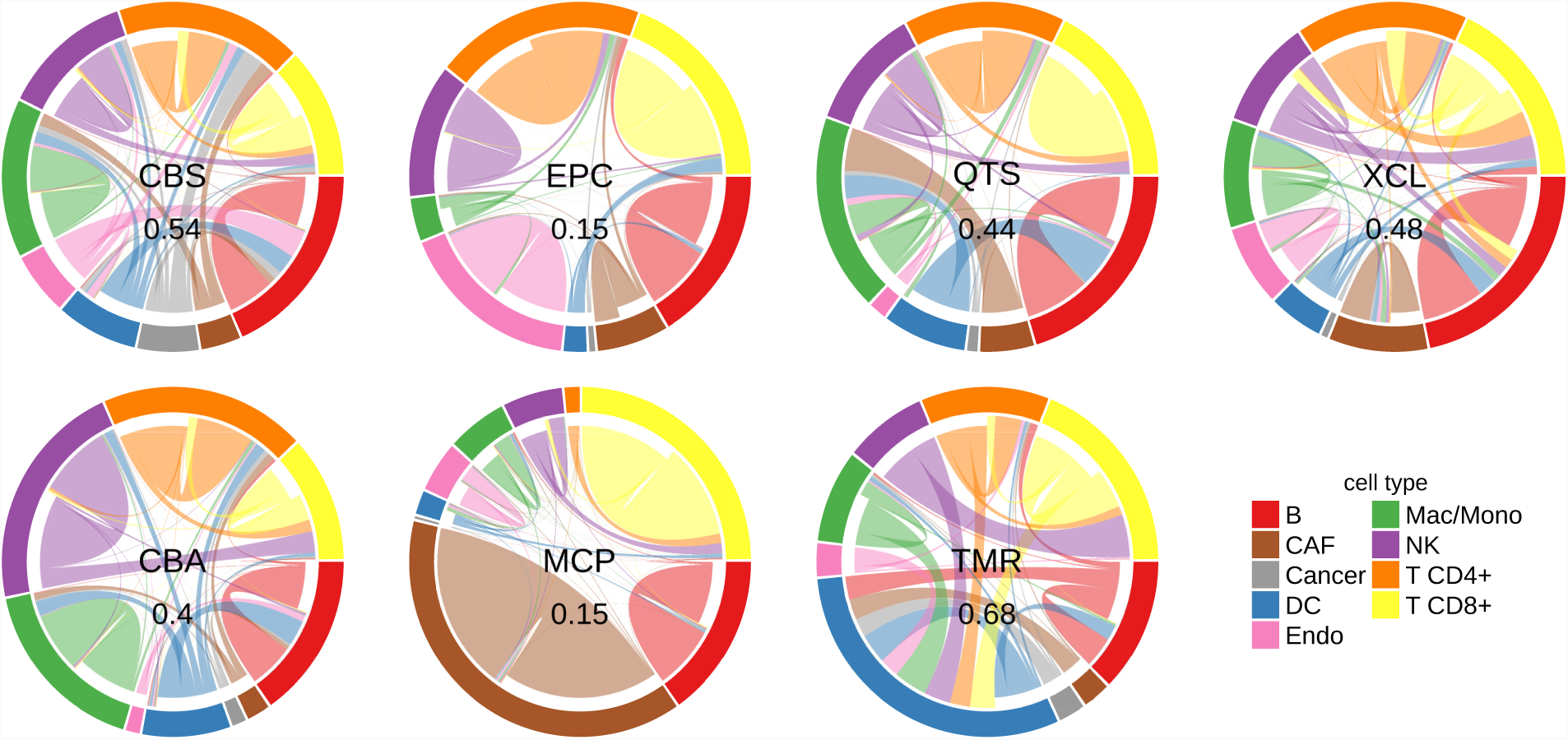
(a) Spillover analysis. All methods were applied to simulated bulk RNA-seq samples containing only cells of one of the nine immune and non-immune cell types. The outer circle indicates the different samples, the connections within refer to the methods’ predictions. The size of a border segment is reflective of the predicted score on that cell type. A connection leading to a border segment of the same color indicates a correctly predicted cell type fraction; a connection leading to a different color indicates spillover, i.e. a prediction of a different cell type than actually present. Note that not all methods provide signatures for all cell types, in that case the connections are indicative of the cell types wrongly predicted when a method is confronted with cell types it has not been optimized for. CD4+ T cell samples are an aggregate of regulatory and non-regulatory CD4+ T cells. The numbers in the center indicate the overall noise ratio, i.e. the fraction of predictions that are attributed to a wrong cell type. Method abbreviations: table 1.

In the previous section, we observed, for instance, that quanTIseq is affected by a high background prediction level for macrophages/-monocytes (figure 2). In the spillover analysis in figure 3, we note that quanTIseq predicts a pure CAF sample to contain a considerable amount of macrophages/monocytes. We, therefore, suspect that the high background prediction level is driven by non-specific marker genes in the quanTIseq signature matrix. Indeed, we identified five genes, *CXCL2, ICAM1, PLTP, SERPING1* and *CXCL3* that are expressed in both CAFs and Macrophages/Monocytes. After removing these genes from the matrix, the background prediction level is significantly reduced by 27 % (figure 4a).

**Figure 4.**
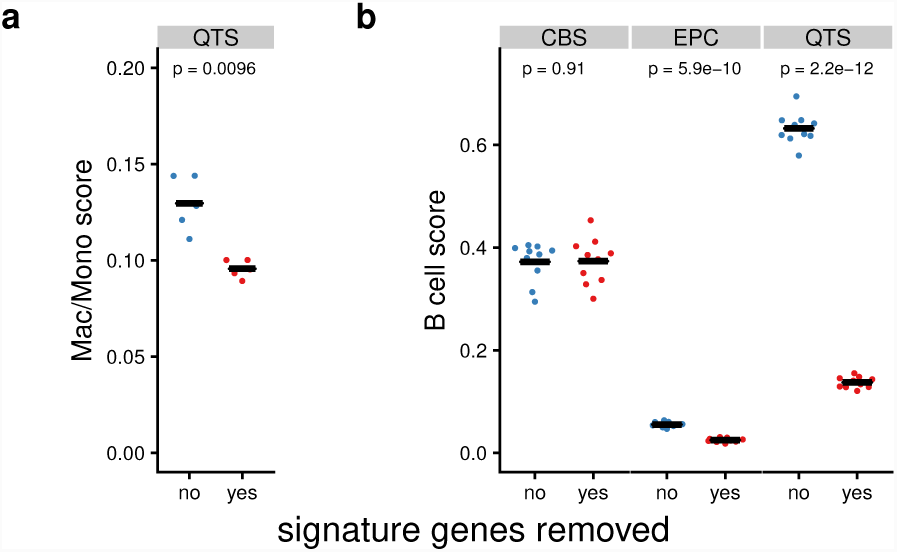
(a) Background prediction level of quanTIseq before and after removing non-specific signature genes. This plot is based on the same five simulated samples used to determine the background prediction level in the Mac/Mono panel of figure 2. (b) B cell score on ten simulated pDC samples before and after removing nonspecific signature genes. Method abbreviations: table 1.

Beyond, for all methods, we consistently observe spillover between CD8+ and CD4+ T cells, between NK cells and CD8+ T cells and from DCs to B cells. The former two spillover effects are conserved in the validation dataset (supplementary figure 6) supporting that the effects are not due to misannotated cells in the single cell dataset.

The spillover between DCs and B cells could not be confirmed in the validation dataset, however, we demonstrate that in the single cell dataset, the B cell and DC clusters are both distinct and well-annotated (supplementary figure 7). Proceeding analogously to the CAF/macrophage spillover above, we identified six genes, *TCL1A, TCF4, CD37, SPIB, IRF8*, and *BCL11A*, that are expressed both in the B cell and DC subpopulations on the mRNA level. Indeed, in the LifeMap Discovery database (Edgar *et al.* (2013)), all six genes are annotated as being expressed in both B cells and plasmacytoid dendritic cells (pDCs). When excluding these genes from the deconvolution matrices, the spillover effect is substantially reduced by 55 % for EPIC and 78 % for quanTIseq (figure 4b). Given that the single cell dataset contains pDCs, quanTIseq has been trained on myeloid dendritic cells (mDCs) and EPIC does not consider any kind of DCs (supplementary table 2), it is not surprising that both methods view these genes as B cell markers and are not able to discriminate between pDCs and B cells.

### Guidelines for method selection

In table 2, we provide guidelines on which method to use for which cell type based on three criteria: interpretability of the score, overall performance, and possible limitations. It is very important to understand the implications of the different scoring strategies. EPIC and quanTIseq are the only methods providing an ‘absolute score’ that represents a cell fraction. All other methods provide scores in arbitrary units, that are only meaningful in relation to another sample of the same dataset. For this reason, and due to a robust overall performance, we recommend EPIC and quanTIseq for general purpose deconvolution. In practice, absolute scores are not always necessary. For instance, in a clinical trial, relative scoring methods can be used to infer fold changes between treatment and control group or to monitor changes in the immune composition in longitudinal samples. In that case, using MCP-counter is a good choice thanks to its highly specific signatures that excelled in the spillover benchmark. A limitation of deconvolution methods is that they are susceptible to background predictions, i.e. prediction of (small) fractions for cell types that are actually absent. Therefore, when interested in presence/absence of a cell type, we suggest using xCell.

**Table 2.**
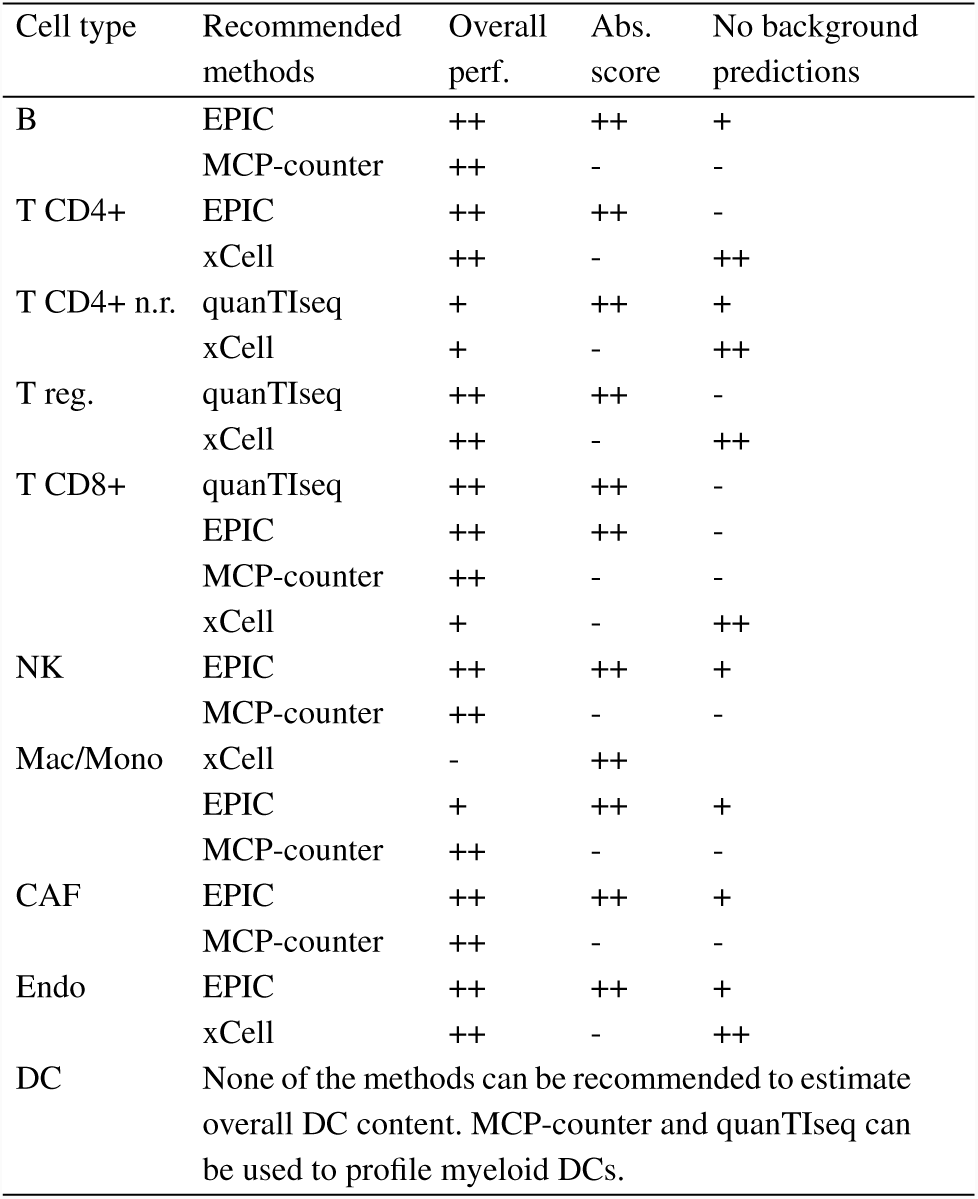
Guidelines for method selection. For each cell type, we recommend which method to use, highlighting advantages and possible limitations. Overall performance: indicates how well predicted fractions correspond to known fractions in the benchmark. Absolute score: the method provides an absolute score that can be interpreted as a cell fraction. Methods that do not provide an absolute score only support inter-sample comparisons within the same experimental dataset, i.e. the score is only meaningful in relation to another sample. Background predictions: indicates, if a method tends to predict a cell type although it is absent.

### Immunodeconv R package provides easy access to deconvolution methods

We created an R package, `immunedeconv`, that provides a unified interface to the different deconvolution methods. The package is freely available from GitHub: https://github.com/grst/immunedeconv. We emphasize the reproducibility and extensibility of our benchmark. A snake-make pipeline (Koster and Rahmann (2012)), including a step-by-step instruction on how to benchmark an additional method is available from https://github.com/grst/immune_deconvolution_benchmark.

## 3 Discussion

Quantification of the cellular composition of bulk tumor samples is a challenging computational problem that has been addressed by a variety of methods. For the utility of these methods, it is imperative to fully understand their capabilities and limitations. Here we provide a comprehensive overview and the first cell type-specific quantitative comparison of existing methods applied to RNA-seq data to guide researchers and clinicians in selecting the most suitable approach for the analysis of their samples.

We assessed seven state-of-the-art computational methods by applying them to simulated bulk RNA-seq samples derived from a scRNA-seq dataset of ∼11,000 single cells, integrated from different experiments (Schelker *et al.* (2017)). We validated the results using true bulk RNA-seq samples profiled by FACS from three independent datasets (Schelker *et al.* (2017); Racle *et al.* (2017); Hoek *et al.* (2015)). FACS or IHC measurements are currently considered the gold standard for assessing cell type composition but in public databases only a few RNA-seq samples with matched FACS or IHC reference can be found. Further publicly-available matched datasets are needed to enhance our ability to reliably compare computational cell-type quantification methods to gold standard techniques. However, even with the limited number of observations available, we showed that there is generally a good agreement between FACS and the methods’ estimates.

ScRNA-seq-based simulation by *in silico* mixing of cell types is a powerful surrogate for bulk-RNA. Not only does it allow for creating an arbitrary number of bulk RNA-seq samples with known cell-type composition, it also uniquely enables us to generate data *ad hoc* to assess specific performance metrics, such as the background prediction level, systematically.

Possible limitations of the single cell-based simulations are related to the transcriptional activities of the cells. First, different cell types contain different amounts of mRNA, differently contributing to bulk RNA-seq samples, while in our simulation approach (averaging normalized gene expression per gene), each cell contributes the same. This would likely severely impact the absolute quantification of cell types, why we refrain from providing an estimate of the absolute deviation for quanTIseq and EPIC. The evaluations performed in this study only rely on between-sample comparisons, and, therefore, should only be marginally affected by this issue, as relative changes are preserved, independent of how much a cell contributes to the mixture. Second, there might be changes in the expression profile of a cell depending on the other cancer-, immune- or stromal cells present in the microenvironment that cannot be taken into account using the simulation approach. A step towards addressing this question could be to evaluate how immune cell profiles depend on the tumor type (see below).

The major conclusion of our study is that RNA-seq can be utilized for estimation of immune cell infiltrates in the tumor robustly and with good accuracy, particularly when a cell type is well characterized. Most methods show near-perfect correlations on CD8+ T-cells, B cells, NK, fibroblasts, and endothelial cells. However, as observed previously by Schelker *et al.* (2017) and Racle *et al.* (2017), regulatory and non-regulatory CD4+ T cells are hard to distinguish. We found that DCs are another example that is difficult to characterize based on the currently available gene signatures. DCs have a variety of subtypes that feature different expression profiles (Collin *et al.* (2013); Villani *et al.* (2017)). While MCP-counter specifically addresses myeloid dendritic cells (mDCs), all other methods provide general signatures for DCs. However, our results demonstrate that none of the methods actually quantify total DCs, but only specific DC subtypes as reported in supplementary table 2. For these “problematic” cell (sub)types, the development of good signatures is essential to improve deconvolution performance and should be the focus of future research.

Our analysis reveals substantial spillover between DCs and B cells in several methods caused by a subset of marker genes characterized by lower cell-specificity. We validated in a public knowledge base that these genes are indeed markers expressed in both pDCs and B cells on the mRNA level and could significantly reduce the spillover effect by removing the genes from the signature matrices. Similarly, we could substantially reduce the background prediction level of quanTIseq on macrophages/monocytes by excluding non-specific genes. By leveraging recently published large-scale single-cell RNA-seq datasets (Sade-Feldman *et al.* (2018); Lambrechts *et al.* (2018); Azizi *et al.* (2018); Zheng *et al.* (2017); Guo *et al.* (2018)), we envision that our understanding of immune cell types and states can be greatly improved and by explicitly addressing spillover effects, more specific cell type signatures can be distilled.

An open question is if profiles derived from one tumor type also apply to another. Schelker *et al.* (2017), to a certain extent, addressed this question, suggesting that cancer-type specific signatures are required. However, their analysis is limited to CIBERSORT and two tumor types (melanoma and ovarian cancer). In our benchmark, other methods already achieve near-perfect correlations on some cell-types without requiring cancer-specific signatures, suggesting that universal cell-type signatures are possible. As a next step, recently published single-cell datasets of various cancer types (Sade-Feldman *et al.* (2018); Lambrechts *et al.* (2018); Azizi *et al.* (2018); Zheng *et al.* (2017); Guo *et al.* (2018)) could be combined with our pipeline to address this question comprehensively.

## 4 Conclusion

In summary, our results demonstrate that computational deconvolution is feasible at high accuracy, given well-defined, high-quality signatures. These findings indicate that future efforts should prioritize defining cell populations and finding reliable signatures over developing new deconvolution approaches. This study establishes a roadmap to evaluate computational cell-type deconvolution methods and provides guidelines to researchers working in immuno-oncology in selecting appropriate methods for characterization of cellular context of the tumors.

## 5 Methods

### Single-cell data

The dataset of 11,759 single cells from Schelker *et al.* (2017) was obtained through their figshare repository (https://figshare.com/s/711d3fb2bd3288c8483a). We reproduced their analysis using their MATLAB source code and exported the final dataset to continue our analysis in R. We excluded all cells that were classified as “Unknown” from the downstream analysis. Moreover, we did not consider cells originating from PBMC as we were interested in the methods’ performance on cancer samples. Gene expression values are expressed as transcripts per million (TPM).

### Immune-cell reference data

Immune cell reference profiles were obtained from the sources described in supplementary table 3. FASTQ files were extracted from SRA files using the `fastq-dump` command of the SRA toolkit (Leinonen *et al.* (2011)). Reads were aligned and gene expression estimated as TPM using STAR (Dobin *et al.* (2013)) and RSEM (Li and Dewey (2011)) using the `rsem-calculate-expression` command in paired-end mode.

### Validation data

We obtained preprocessed bulk RNA-seq and FACS data for 8 PBMC samples from Hoek *et al.* (2015) through personal communication from the quanTIseq authors (Finotello *et al.* (2017)). We obtained bulk RNA-seq and FACS data for 4 metastatic melanoma samples from Racle *et al.* (2017) through GEO accession number GSE93722. We obtained bulk RNA-seq and FACS data for 3 ovarian cancer ascites samples (Schelker *et al.* (2017)) from the figshare repository (https://figshare.com/s/711d3fb2bd3288c8483a). The 3 ovarian cancer ascites samples consist of two technical replicates each. The two replicates are highly consistent for each of the samples (Pearson’s *r ≥* 0.98), we, therefore, took the straightforward approach of merging them by the arithmetic mean for each gene. All gene expression values are expressed as TPM.

### Cell type mapping

For comparing the methods, it is essential to map the cell types of the different methods to a controlled vocabulary (CV). It has to be taken into account, that the different methods resolve the cell types at different granularities. E.g. while CIBERSORT predicts naive- and memory CD8+ T cells, all other methods only predict CD8+ T cells. We address this issue by creating a hierarchy of cell types, and map each cell type from a dataset or method to a node in the cell type tree (see supplementary figure 8 and supplementary table 4). If a node (e.g. CD8+ T cell) is to be compared among the different methods, the estimated fraction is computed as follows: (1) if the method provides an estimate for the node, take that value. (2) If the method provides an estimate for all child nodes, sum up all children. This is done recursively until no estimates are available or the leaves of the tree are reached. (3) If an estimate is missing for at least one of the child nodes, no estimate will be available. An exception was made for the following cell types, that are only provided by few methods: T cell gamma delta, Macrophage M0, Monocyte, T cell follicular helper. If an estimate is missing for one of those cell types, the remaining child nodes will be summed up.

### Validation of the methodology

Schelker *et al.* (2017) provide three ovarian cancer ascites samples for which both single cell RNA-seq and bulk RNA-seq data is available. We generated artificial bulk samples by taking the mean of the TPM values for each gene of all single-cell samples. First, we compared the values using Pearson correlation on log-scale. Next, we used BioQC (Zhang *et al.* (2017)) to test for differential enrichment of gene ontology (GO)-terms (The Gene Ontology Consortium (2017)). Finally, we ran the deconvolution methods on both the true and simulated bulk RNA-seq sample and compared the results. To demonstrate that xCell’s low correlation is in fact due to little variance between the samples, we reran xCell while additionally including the 51 immune cell reference samples (supplementary table 3) in the expression matrix of both simulated and true bulk RNA-seq.

### Deconvolution methods

We implemented an R package, `immunedeconv`, that provides a unified interface to all seven deconvolution methods compared in this paper. The CIBERSORT R source code was obtained from their website on 2018-03-26. The xCell, EPIC, MCP-counter, TIMER and quanTIseq source codes were obtained from GitHub from the following commits: dviraran/xCell@870ddc39, GfellerLab/EPIC@e5ae8803, ebecht/MCP-counter@e7c379b4, hanfeisun/TIMER@a030bac73, FFinotello/quan-TIseq@ee9f4036.

We ran CIBERSORT with disabled quantile normalization, as recommended on their website for RNA-seq data. While quanTIseq provides an entire pipeline, starting with read-mapping and estimation of gene expression, we only ran the last part of that pipeline, which estimates the immune cell fractions from gene expression data. We ran TIMER with “OV” on ovarian cancer ascites samples and with “SKCM” on melanoma samples. We ran quanTIseq with the option `tumor = TRUE` on all tumor samples and `tumor = FALSE` on the PBMC samples. We ran EPIC with the `TRef` signature set on all tumor samples and with `BRef` on the PBMC samples. We ran xCell with the `cell.types.use` parameter to avoid overcompensation by the spillover correction. For simulated tumor data, `cell.types.use` was set to B, CAF, DC, Endo, Mac/Mono, NK, T CD4+ n.r., T CD8+, T reg. For the validation datasets, it was set to B, DC, Mono, NK, T CD4+, T CD8+, T. We disabled the mRNA scaling options of quanTIseq and EPIC for the single cell simulation benchmark using the `mRNAscale` and `mRNA_cell` options, respectively. Notably, this only has an effect on the absolute values, but not on the correlations used to compare all methods. For each of the datasets (simulated, Hoek, Schelker, Racle), we submitted all samples in a single run.

### Simulation benchmark

We simulated bulk RNA-seq samples by calculating the average gene expression per gene of cells sampled from the single cell dataset. Independently for each sample, 500 cells were randomly selected as follows: A fraction *f* of cancer cells was drawn from *N* (0.33, 0.3), which is the empirical distribution of cancer cells in the melanoma and ascites samples in the single cell dataset. *f* was constrained to the interval [0, 0.99]. Half of the samples use melanoma cancer cells, the other half ovarian cancer cells (3) The remaining fraction 1 *- f* was randomly assigned to immune cells, resulting in a fraction vector. (4) The fraction vector was multiplied with 500 to obtain a cell count vector. (5) The corresponding number of cells was drawn with replacement from the single cell dataset. That implies that for cell populations with only a few cells available, the artificial bulk sample will contain the same single cell sample multiple times.

Known fractions of simulated samples are compared to the predicted scores using Pearson’s correlation coefficient *r*. P-values and *r* have been computed using the `cor` function in R.

### Minimal detection fraction and background prediction levels

We simulated bulk RNA-seq samples as follows: For each cell type *c*, we create five independent samples for each *i ∈ {*0, 5, 10, *…,* 50, 60, *…,* 100, 120, *…,* 300, 350, *…,* 500*}*. For each sample independently, we randomly sampled *i* cells of type *c* and a background of 1000 cells containing all cell types except *c* from the single cell dataset. The selected cells were aggregated by calculating the arithmetic mean for each gene. This results in five batches of 300 samples each (30 spike-in levels *×* 10 cell types). Each of the five batches was submitted to the methods independently in a single run.

We quantify the background prediction level of cell type *c* as the predicted score of *c* in samples of type *c* with *i* = 0, *i.e.* in samples with no cells of type *c* present. We quantify the minimal detection fraction as the minimal *i* at which the predicted score of *c* is significantly different from the background prediction level (one-sided t-test, alpha = 0.05).

### Spillover

We refer to spillover as the erroneous prediction of another cell type due to a partial overlap of the signatures. We measured spillover on two datasets: (1) 44 true bulk RNA-seq samples of pure immune cells (supplementary table 3). (2) 45 simulated bulk RNA-seq samples of nine cell types. For each cell type *c*, we simulated five samples by independently aggregating 500 random cells of type *c* by taking the arithmetic mean. Each dataset was submitted to the methods independently in a single run. We visualized the results as chord diagrams (figure 3) using the R package circlize (Gu *et al.* (2014)).

### Other tools

We generated reproducible reports using bookdown 0.7. We use Snakemake 5.2.0 (Koster and Rahmann (2012)), conda and bioconda (Grüning *et al.* (2018)) to integrate our analyses in a reproducible pipeline.

## Supporting information

Supplementary Table 1: GO term enrichment

Supplementary Table 3: Immune cell reference profiles

Supplementary Table 4: Cell type mapping

## Code availability

An R package providing a unified interface to the deconvolution methods is freely available from GitHub: https://github.com/grst/immunedeconv. A ready-to-run Snakemake pipeline, including preprocessed data, to reproduce the results is available from https://github.com/grst/immune_-deconvolution_benchmark.

### Acknowledgements

We would like to thank Hubert Hackl for helpful discussions.

## Supplementary information

Supplementary data are available at *Bioinformatics* online.

