## supplementary material for "Comprehensive evaluation of computational cell-type quantification methods for immuno-oncology"

### Supplementary Figures

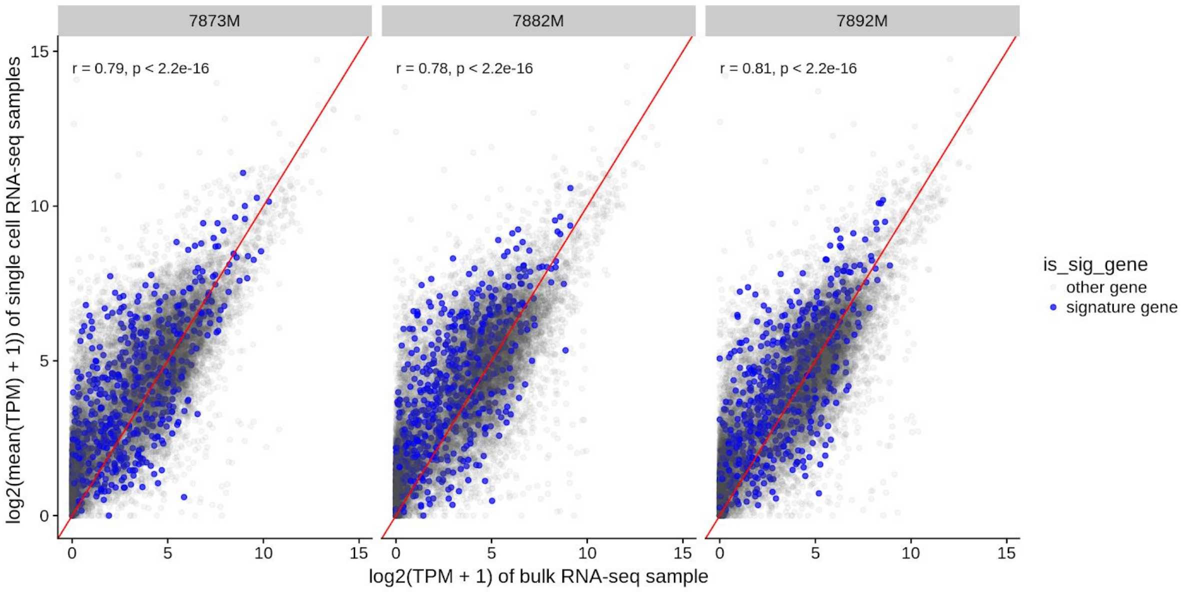


**Supplementary figure 1:** Correlation of simulated bulk samples with corresponding genuine bulk RNA-seq samples. Schelker et al.[1] provide three samples for which paired single cell and bulk RNA-seq has been performed. We generate simulated bulk RNA-seq samples by taking the average of the single cells and compare the simulated to the genuine samples. Genes used in the signatures of quanTIseq, EPIC, CIBERSORT and MCP-counter are shown in blue. *r* indicates Pearson correlation. A perfect correlation cannot be expected, due to differences in single-cell dissociation efficiencies[2].

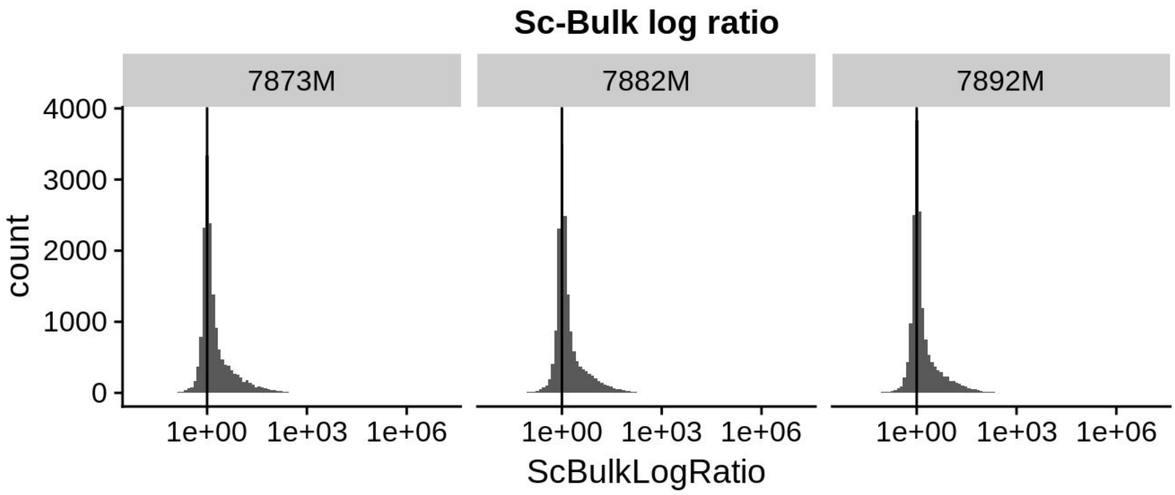


**Supplementary figure 2:** Log-ratio of gene expression between simulated and genuine bulk RNA-seq samples. Schelker et al.[1] provide three samples for which paired single cell and bulk RNA-seq has been performed. We generate simulated bulk RNA-seq samples by taking the average of the single cells and compare the simulated to the genuine samples. We were interested if there is a systematic bias in simulated vs. genuine bulk RNA-seq samples. To this end, we calculate the sc-bulk log ratio by $\log(sc+1)/\log(bulk+1)$. We observe that the distribution is skewed in direction of the single cells, i.e. more genes have increased gene expression in sc-aggregates rather than decreased. In supplementary table 1 we demonstrate that despite of this observation, there is no systematic bias between the two samples as far as immune-related genes are concerned.

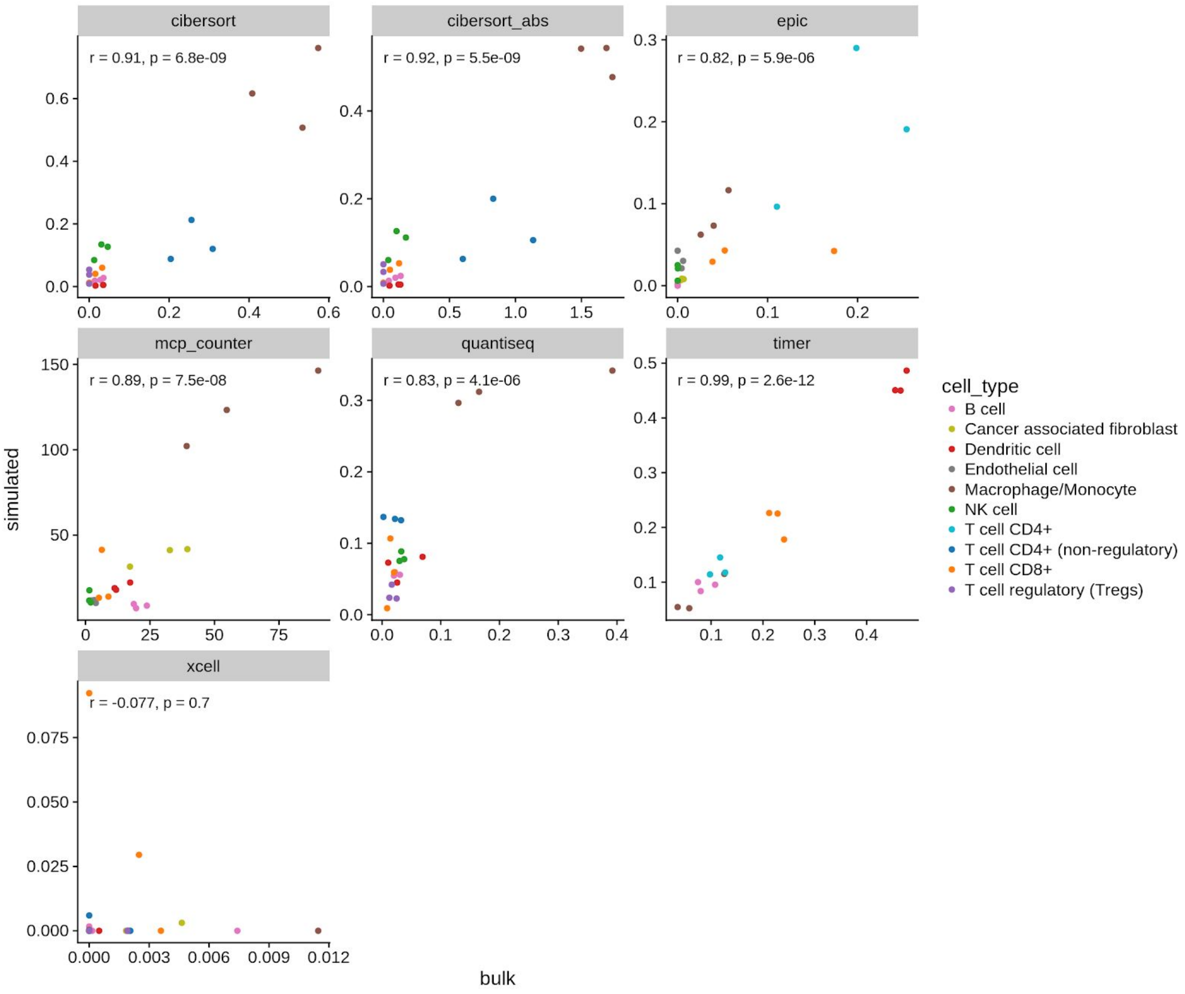


**Supplementary figure 3:** Predictions of the methods on simulated and genuine bulk RNA-seq samples. Schelker et al.[1] provide three samples for which paired single cell and bulk RNA-seq has been performed. We generate simulated bulk RNA-seq samples by taking the average of the single cells. A perfect correlation cannot be expected, due to differences in single-cell dissociation efficiencies[2].

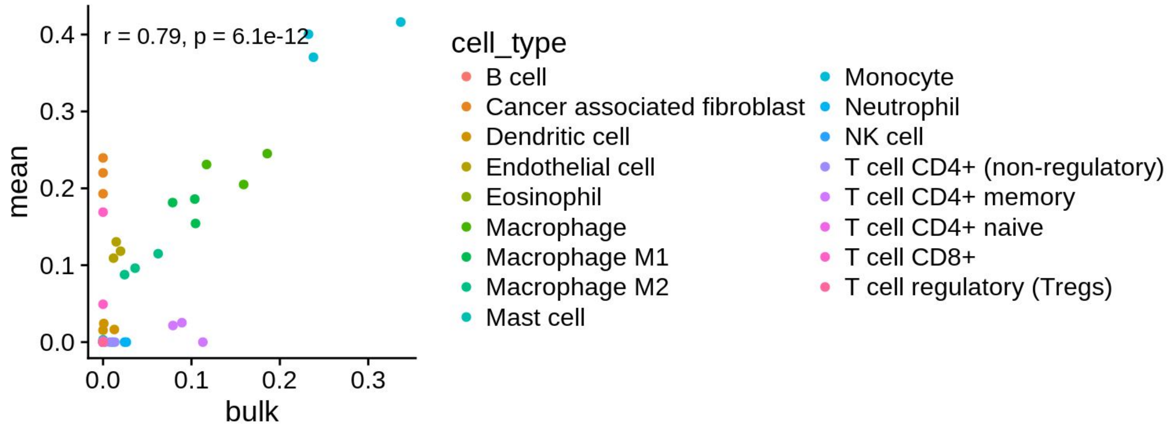


**Supplementary figure 4**: In supplementary figure 2, we observe low correlations and low absolute predictions for xCell. xCell relies on the inter-sample variance to compute scores (see their README on GitHub). Therefore we hypothesized that low inter-sample variance is at the bottom of this observation. We ran xCell on the same three samples, but this time included 51 immune cell reference samples (supplementary table 3) in the same run. This figure shows the correlation of xCell predictions on simulated vs. genuine bulk RNA-seq samples with the 51 additional samples included in the same run.

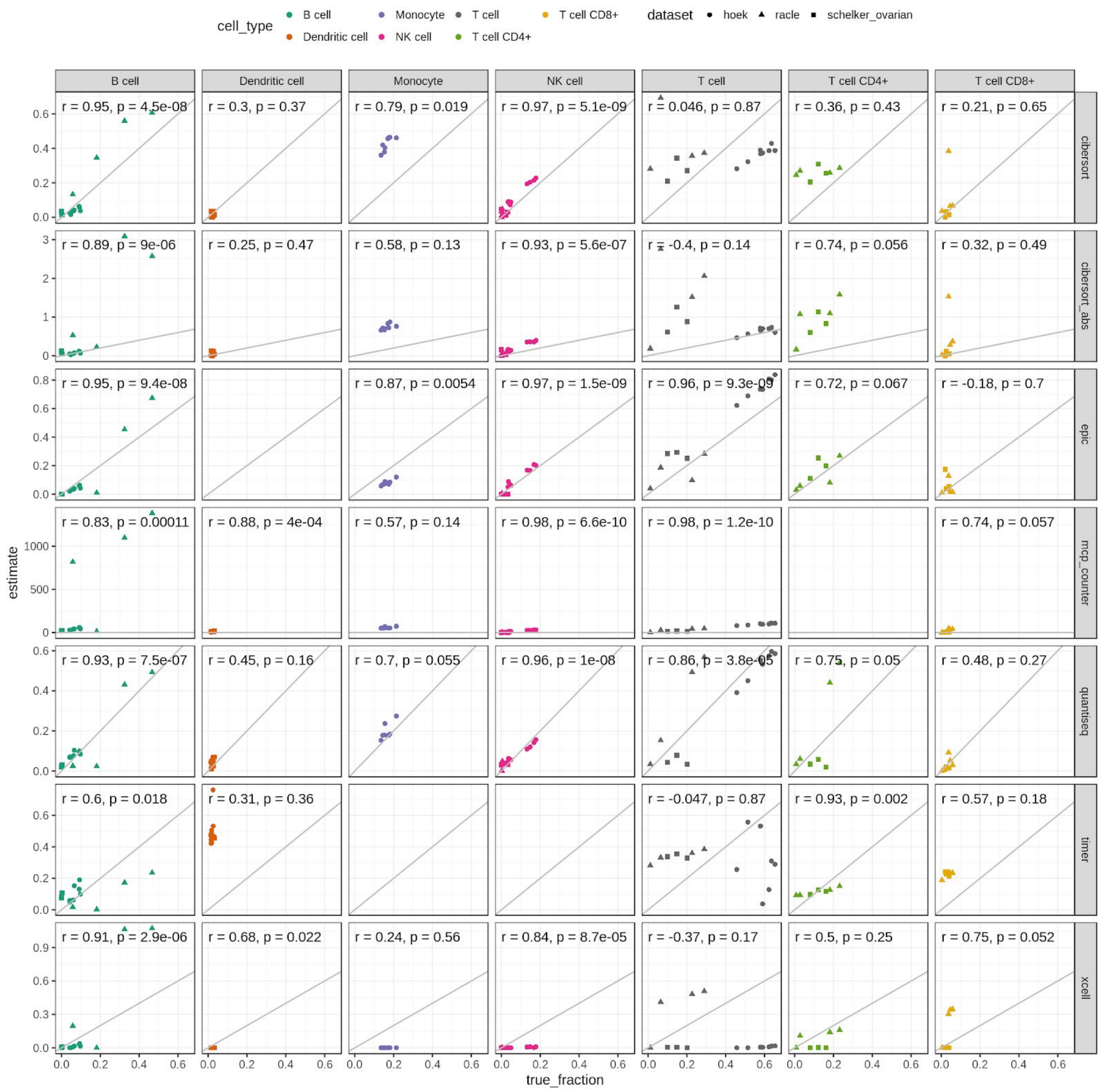


**Supplementary figure 5:** Correlations of known vs. predicted cell type fractions on 15 genuine bulk RNA-seq samples from three datasets[1,3,4] profiled with FACS. Correlations are shown for each cell type independently, to enable a fair comparison for methods that only allow for inter-sample comparisons. Note that for $CD8^+$ T cells (yellow) the correlations are low, but also the inter-sample variance is very low, so that the correlation is not a good measure here.

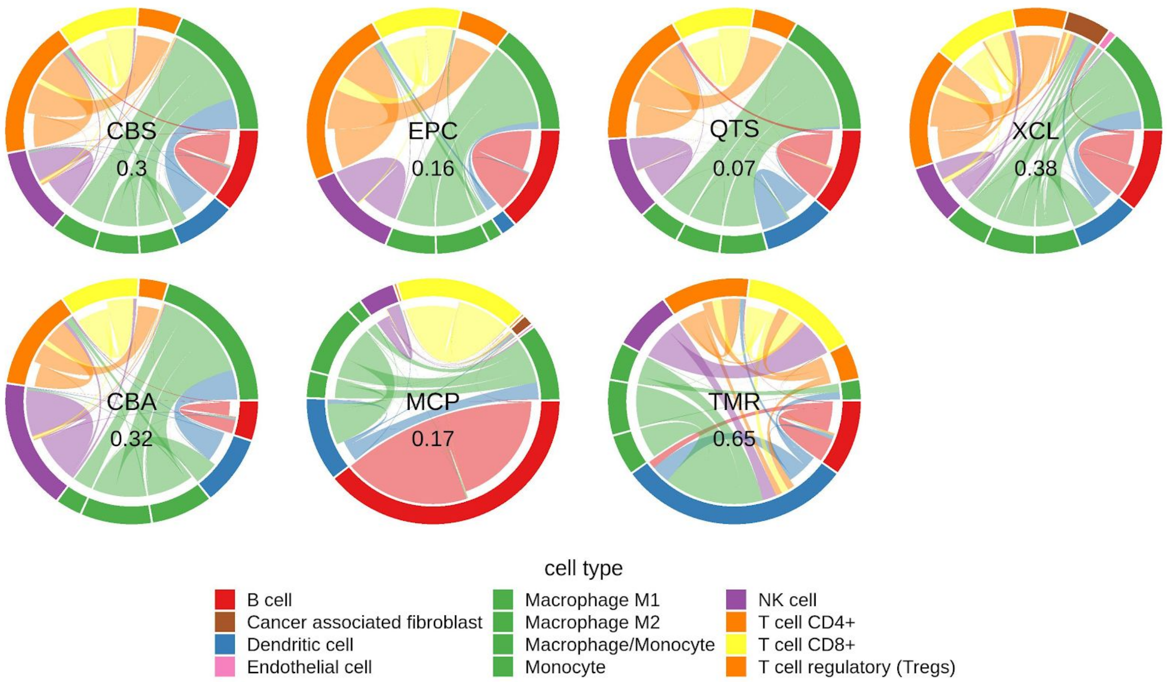


**Supplementary figure 6:** Spillover analysis on validation dataset. All methods were applied to genuine bulk RNA-seq datasets of FACS-purified immune cells. The outer circle indicates the different samples, the connections within refer to the methods' predictions. The size of a border segment is reflective of the predicted score on that cell type. A connection leading to a border segment of the same color indicates a correctly predicted cell type fraction; a connection leading to a different color indicates spillover. The numbers in the center indicate the overall noise ratio, i.e. the fraction of predictions that are attributed to a wrong cell type. Note that the immune reference data has been used to derive signatures for quanTIseq[5]. Its performance on this dataset is therefore likely over-estimated.

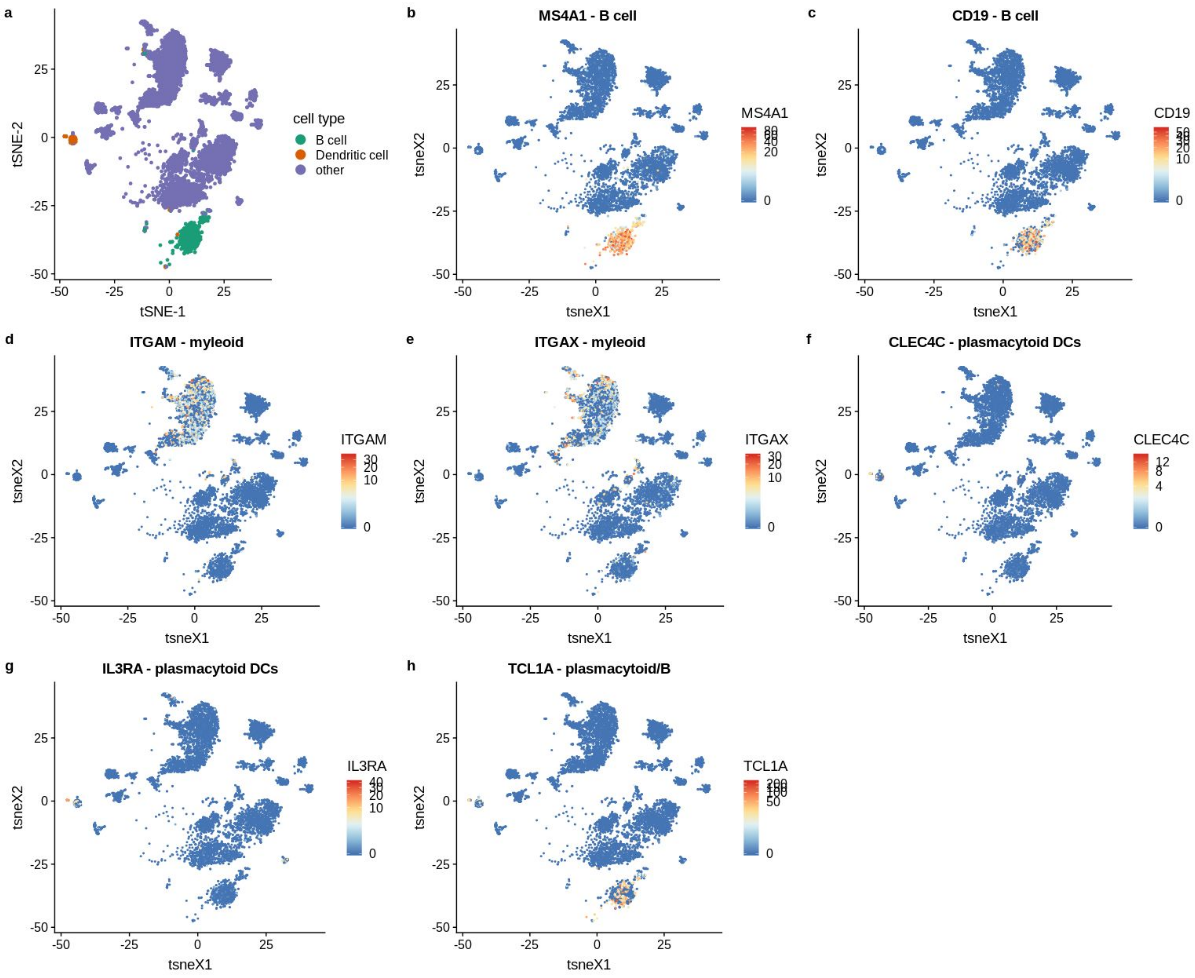


**Supplementary figure 7:** (a) DC and B cell clusters are distinct in the single cell dataset. (a-h) The DC and B cell clusters are well-annotated. The panels show the expression (log TPM) of marker genes[1,6] for B cells, plasmacytoid DCs and myeloid cells across the single cell dataset.

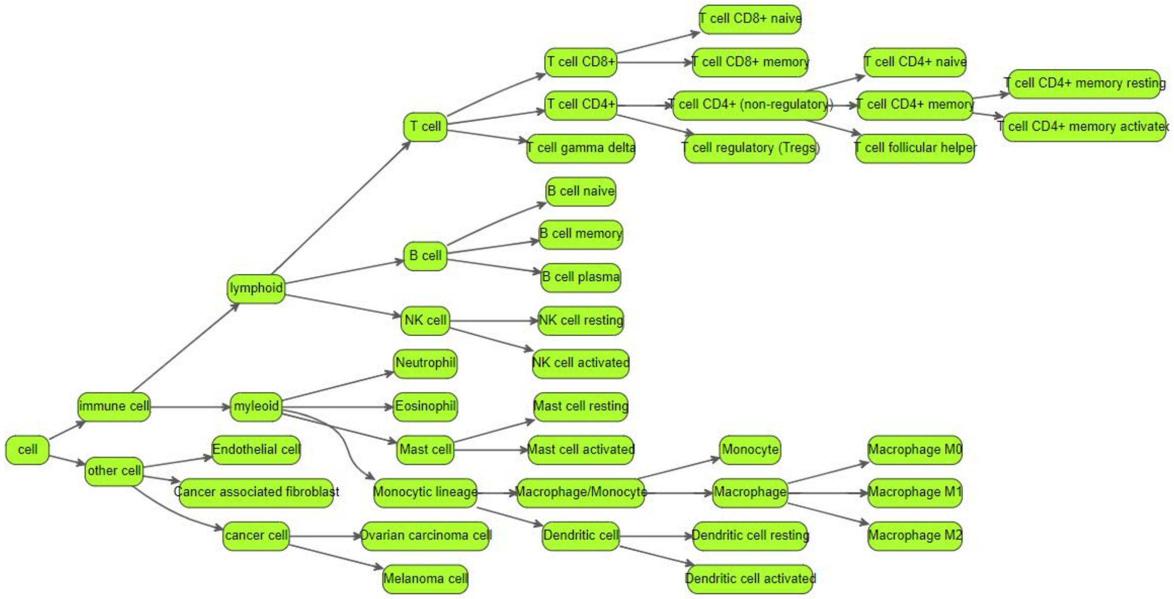


**Supplementary figure 8:** Hierarchy of immune cell types used for mapping cell types between methods and datasets. For comparing the methods, it is essential to map the cell types of the different methods to a controlled vocabulary (CV). It has to be taken into account, that the different methods resolve the cell types at different granularities. E.g. while CIBERSORT predicts naive- and memory $CD8^+$ T cells, all other methods only predict $CD8^+$ T cells. We address this issue by creating a hierarchy of cell types, and map each cell type from a dataset or method to a node in the cell type tree. The mapping of cell types to the hierarchy is detailed in supplementary table 4.

### Supplementary Tables

**Supplementary Table 1:** Gene-ontology (GO)-term enrichment in simulated vs. genuine bulk RNA-seq samples. In supplementary figure 2 we observed that more more genes have increased gene expression in sc-aggregates rather than decreased. We were wondering if there is a systematic bias towards a certain group of genes that tend to be over/underrepresented in single-cell vs. bulk RNA seq. To this end, we ran a gene set enrichment test with BioQC[7] using gene ontology[8] (BP and CC terms) as the knowledge base. **(a)** In this sheet we display terms that, under moderate stringency of filtering (Benjamini-Hochberg FDR < 0.01), shows significant enrichment in either direction. We notice that among these significantly enriched gene-sets above, no gene-sets are directly involved in immune response, cytokine/interleukin/chemokine response. **(b)** In this sheet, we show the enrichment of all GO-terms containing one of the keywords *inflammation, cytokine, chemokine, interleukin, antigen, macrophage, dendritic cell* and found none of the terms to be differentially enriched.

*Table is supplied as separate Excel Sheet.*
[https://www.dropbox.com/s/f4a4ow79aa6s0lu/supplementary%20table%201_%20GO-term%20enrichment.xlsx?dl=0](https://www.dropbox.com/s/f4a4ow79aa6s0lu/supplementary%20table%201_%20GO-term%20enrichment.xlsx?dl=0)

**Supplementary Table 2:** Subtypes of dendritic cells (DC) estimated by the different methods. Except for MCP-counter none of the methods explicitly addresses a certain DC subtype, yet we could reconstruct this information from the methods' supplementary information.

| dataset/method | subtype | reference |
|---|---|---|
| Schelker[1] | plasmacytoid DC | Identified in the single cell data using CD123 and CD303 marker genes[1] which are pDC marker genes according to [6]. |
| Hoek[3] | myeloid DC | primary human myeloid DC according to annotation on GSE64655 |
| MCP-counter[9] | myeloid DC | signature explicitly annotated as myeloid DC |
| CIBERSORT[10] | monocyte-derived DC | "Monocytes isolated as above were cultured in RPMI with 10% heat-inactivated FBS, 1 × Pen/Strep, 2 mM L-glutamine, 10 mM HEPES, 1 mMsodium pyruvate, then differentiated into dendritic cells by 17 ng/ml IL4, and 67 ng/ml GMCSF for 5 days at 5 × 106 cells/ml." (GSE22886)[11] |
| quanTIseq[5] | myeloid DC | signatures derived from Hoek[3] data |
| EPIC[4] | (no DC signature provided) | |
| TIMER[12] | monocyte-derived DC | training data is a mix of various monocyte-derived DCs from HPCA (See table S8 of [12]) |
| xCell[13] | myeloid DC | uses a combination of various, mostly myeloid, DC samples (personal communication with authors) |

**Supplementary Table 3:** List of 51 immune cell reference samples, originally curated by Finotello *et al.*[5].

*Table is supplied as separate Excel Sheet.*
https://www.dropbox.com/s/75cmtlbjv6msmg8/supplementary%20table%203_%20immune%20cell%20reference%20data.xlsx?dl=0

**Supplementary Table 4:** Cell type mapping. Different methods use different terms and resolve cell types at different resolution. **(a)** In this sheet, we define a controlled vocabulary organized as a hierarchy of cell types. **(b)** In this sheet, we mapped cell types from each dataset and method to a node of the cell type tree defined in (a).

*Table is supplied as separate Excel Sheet.*
https://www.dropbox.com/s/hyt9j0xd63d55xi/supplementary%20table%204_%20cell%20type%20mapping%20.xlsx?dl=0
